# Extratubular polymerized uromodulin induces leukocyte recruitment and inflammation *in vivo*

**DOI:** 10.1101/2020.07.18.206508

**Authors:** Roland Immler, Bärbel Lange-Sperandio, Tobias Steffen, Heike Beck, Jonas Roth, Georg Hupel, Frederik Pfister, Bastian Popper, Bernd Uhl, Hanna Mannell, Christoph A. Reichel, Volker Vielhauer, Jürgen Scherberich, Markus Sperandio, Monika Pruenster

## Abstract

Uromodulin (UMOD) is produced and secreted by tubular epithelial cells. Secreted UMOD polymerizes (pUMOD) within the lumen, where it regulates salt transport and protects the kidney from bacteria and stone formation. Under various pathological conditions, pUMOD accumulates within the tubular lumen and reaches extratubular sites where it may interact with renal interstitial cells. Here, we investigated the potential of extratubular pUMOD to act as a damage associated molecular pattern (DAMP) molecule thereby creating local inflammation. We found that intrascrotal and intraperitoneal injection of pUMOD induced leukocyte recruitment *in vivo* and led to TNF-α secretion by F4/80 positive macrophages. Additionally, pUMOD directly affected vascular permeability and increased neutrophil extravasation independent of macrophage-released TNF-α. Interestingly, pUMOD did not directly upregulate adhesion molecules on endothelial cells and did not directly activate β2 integrins on neutrophils. In obstructed neonatal murine kidneys, we observed extratubular UMOD accumulation with tubular atrophy and leukocyte infiltrates. Finally, we found extratubular UMOD deposits associated with peritubular leukocyte infiltration in kidneys from patients with inflammatory kidney diseases. Taken together, we identified extratubular pUMOD as a strong inducer of leukocyte recruitment, underlining its critical role in mounting an inflammatory response in various kidneys pathologies.

## INTRODUCTION

Uromodulin (UMOD), also known as Tamm-Horsfall protein (THP), is a glycoprotein with a molecular weight of 80-90kDa expressed by epithelial cells lining the thick ascending limb (TAL) of the loop of Henle and to minor degree the early distal convoluted tubules (1, 2). Under physiological conditions, the glycosylphosphatidylinositol (GPI) anchored protein is located on the apical plasma membrane (3), where it is released into the tubular lumen via conserved proteolytic cleavage (4). Cleaved UMOD polymerizes (pUMOD) and represents the most abundant urinary protein in healthy humans, with a secretion rate of 50-150mg/day (2). Within the lumen, pUMOD has numerous functions. It contributes to the water impermeability of the TAL and affects ion transport by binding to the Na^+^-K^+^-2Cl^-^ cotransporter NKCC2 (5) and the potassium channel ROMK (6), hence regulating renal function and blood pressure (7). Furthermore, pUMOD blocks binding of *E*.*coli* to uroplakin on the urothelium (8) and protects from urinary tract infections (9, 10). The polymerized and negatively charged structure of the protein inhibits calcium crystal creation, prevents kidney stone formation (11, 12) and regulates renal magnesium homeostasis (13). A smaller fraction of UMOD is also released basolaterally into the interstitium and a monomeric form of UMOD (mUMOD) is detectable in the serum (14). The function of serum mUMOD is still a matter of debate. Reduced serum levels correlate with higher cardiac mortality (15), elevated systemic ROS (16) and increased susceptibility to develop type 2 diabetes mellitus accompanied by reduced glucose metabolism (17). In addition, low serum levels were shown to be associated with reduced GFR (18), linking mUMOD serum levels to kidney function and survival. Abnormal distribution of UMOD, intracellular accumulation within the ER and the cytoplasm occurs in patients with autosomal dominant tubulointerstitial kidney disease (ADTKD) (19-22). ADTKDs are caused by mutations in the genes encoding for renin, mucin 1 or uromodulin (23, 24) and are characterized by tubulointerstitial fibrosis, tubular atrophy and chronic renal failure (22). In patients with multiple myeloma, pUMOD co-precipitates with monoclonal immunoglobulin free light chains and forms casts which block urinary flow and cause tubular atrophy resulting in progressive interstitial inflammation and fibrosis (25). Finally, in mouse models of unilateral ureteral obstruction (UUO) accompanied with an inflammatory response, pUMOD accumulation and extratubular deposition has been documented (26, 27). These findings imply a potential proinflammatory role of pUMOD under pathological conditions. However, direct *in vivo* evidence to support this function is still missing. To clarify a putative proinflammatory role of pUMOD and differentiate pUMOD effects from the immunomodulatory role already described for mUMOD, we investigated leukocyte recruitment in mouse models for microvascular inflammation. We performed intravital microscopy in the cremaster muscle and found a marked induction of leukocyte adhesion and extravasation following intrascrotal injection of pUMOD. Intraperitoneal application of pUMOD induced recruitment of neutrophils and inflammatory monocytes as determined by flow cytometry. We also identified an upregulation and abnormal localization of UMOD accompanied by leukocyte infiltration in a mouse model of neonatal obstructive nephropathy and in renal biopsies from patients with various chronic renal inflammatory disorders. Taken together, our study expands the current knowledge of the broad spectrum of UMOD functions, proposing a proinflammatory and damage associated molecular pattern (DAMP)-like role of extratubular pUMOD in kidney pathologies.

## RESULTS

### pUMOD induces leukocyte recruitment in murine microvascular inflammation models

To study potential proinflammatory properties of pUMOD (for the polymerized structure see **Supplemental Figure 1A**) and its role in leukocyte recruitment *in vivo*, we applied an intravital microscopy model of acute microvascular inflammation in the mouse cremaster muscle. We injected pUMOD isolated from human urine into the scrotum (i.s.) of WT mice. Two hours after pUMOD injection, we exteriorized the cremaster muscle and assessed leukocyte rolling flux fraction, number of adherent leukocytes, and leukocyte rolling velocities in postcapillary cremaster muscle venules (28). Application of pUMOD led to a decrease in leukocyte rolling flux fraction and an increase in the number of adherent leukocytes/mm^2^ compared to i.s. injection of normal saline (control) or human albumin solution (hALB), indicating a proinflammatory role of injected pUMOD (**Figure 1A, 1B and Supplemental Movie 1 and 2**). Of note, hemodynamic parameters (vessel diameter, blood flow velocity, shear rate, and systemic white blood cell count) did not differ between the groups (**Supplemental Table 1**). Further, administration of pUMOD reduced rolling velocities compared to injection of normal saline (**Figure 1C**), corroborating the assumed induction of inflammation by pUMOD. Severe local inflammation goes along with an upregulation of E-selectin on the affected vascular endothelium (29). To functionally test for E-selectin expression on the inflamed endothelium, we applied an E-selectin blocking antibody, which significantly increased leukocyte rolling velocity in pUMOD treated mice (**Figure 1C**) demonstrating endothelial expression of E-selectin.

**Figure 1.**
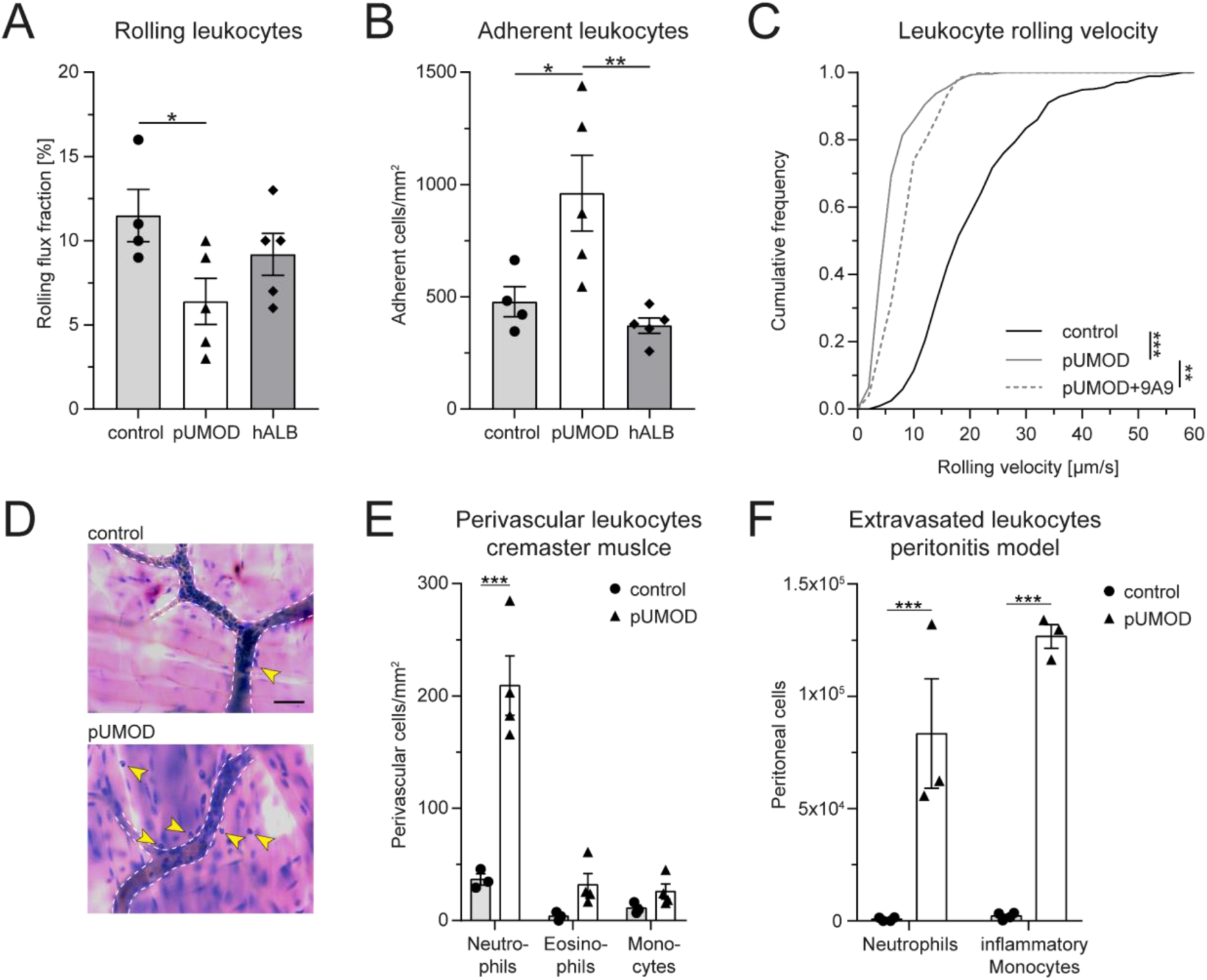
pUMOD induces leukocyte recruitment in murine microvascular inflammation models. pUMOD, normal saline (control) or human albumin (hAlb) were injected i.s. two hours prior to intravital microscopy of postcapillary venules of the mouse cremaster muscle. (**A**) Leukocyte rolling flux fraction and (**B**) number of adherent leukocytes in 17 (control), 23 (pUMOD) and 18 (hALB) venules of n=4-5 mice per group were assessed in postcapillary venules of mouse cremaster muscles. (**C**) Leukocyte rolling velocities of n=392 (control), 274 (pUMOD) and 123 (pUMOD+9A9) cells in 4-5 mice per group were analyzed. pUMOD administration reduced rolling flux fraction and rolling velocity of leukocytes and increased the number of adherent cells compared to control conditions (one-way ANOVA, Tukey’s multiple comparison). (**D**) Giemsa staining (representative micrographs of n=3-4 mice per group) of cremasteric tissue were carried out. Arrow bars indicate perivascular leukocytes (scale bar=30µm). (**E**) Number of perivascular neutrophils, eosinophils and monocytes were calculated in 37 (control) and 54 (pUMOD) perivascular regions of n=3-4 mice per group (two-way repeated measurements ANOVA, Sidak’s multiple comparison). (**F**) pUMOD and normal saline (control) was injected i.p. and numbers of recruited neutrophils and inflammatory monocytes into the peritoneum was assessed 6 hours later (n=3-4 mice per group, two-way ANOVA, Sidak’s multiple comparison). *: p≤0.05, **: p≤0.01, ***: p≤0.005, data is presented as mean±SEM, cumulative frequency or representative images.

Next, we aimed to study the effect of pUMOD on leukocyte extravasation. To do this, we injected pUMOD into the scrotum of WT mice. Two hours after injection, we removed and fixed the cremaster muscles before staining the tissue with Giemsa (**Figure 1D**) in order to analyze the number of perivascular neutrophils, eosinophils, and monocytes. In contrast to normal saline injection, pUMOD application induced extravasation of neutrophils within 2h (**Figure 1E**). Neutrophils are the first cells reaching sites of inflammation, while monocytes accumulate at later time points and persist longer. In order to test the putative capability of pUMOD to induce monocyte recruitment as well, we applied pUMOD into the peritoneum of WT mice and assessed leukocyte recruitment 6h after injection of pUMOD (**Figure 1F**). While application of normal saline did not induce any leukocyte extravasation, intraperitoneal stimulation with pUMOD led to substantial recruitment of neutrophils and monocytes into the inflamed peritoneal cavity.

To demonstrate that murine polymerized UMOD (murine pUMOD) is also able to induce inflammation in the mouse cremaster muscle, we additionally injected a recombinant form of the mouse protein i.s. This truncated form of murine UMOD (aa25-588) contains all important domains of UMOD (30) and also forms polymers (**Supplemental Figure 1B**). Similar to our findings with human pUMOD, murine pUMOD reduced leukocyte rolling flux fraction and increased the number of adherent cells (**Supplemental Figure 2A** and **B**) compared to the injection of mouse albumin (mALB). Rolling velocities of leukocytes were similar in mice treated with pUMOD and murine pUMOD (**Supplemental Figure 2C**). Again, hemodynamic parameters did not differ between the groups (**Supplemental Table 2**). We also analyzed the effect of murine pUMOD on leukocyte extravasation. Similar to pUMOD, injection of murine pUMOD induced extravasation of primarily neutrophils (**Supplemental Figure 2D**). Taken together, these data clearly illustrate a proinflammatory function of pUMOD *in vivo*, affecting leukocyte rolling, leukocyte rolling velocity, leukocyte adhesion, and leukocyte extravasation.

**Figure 2.**
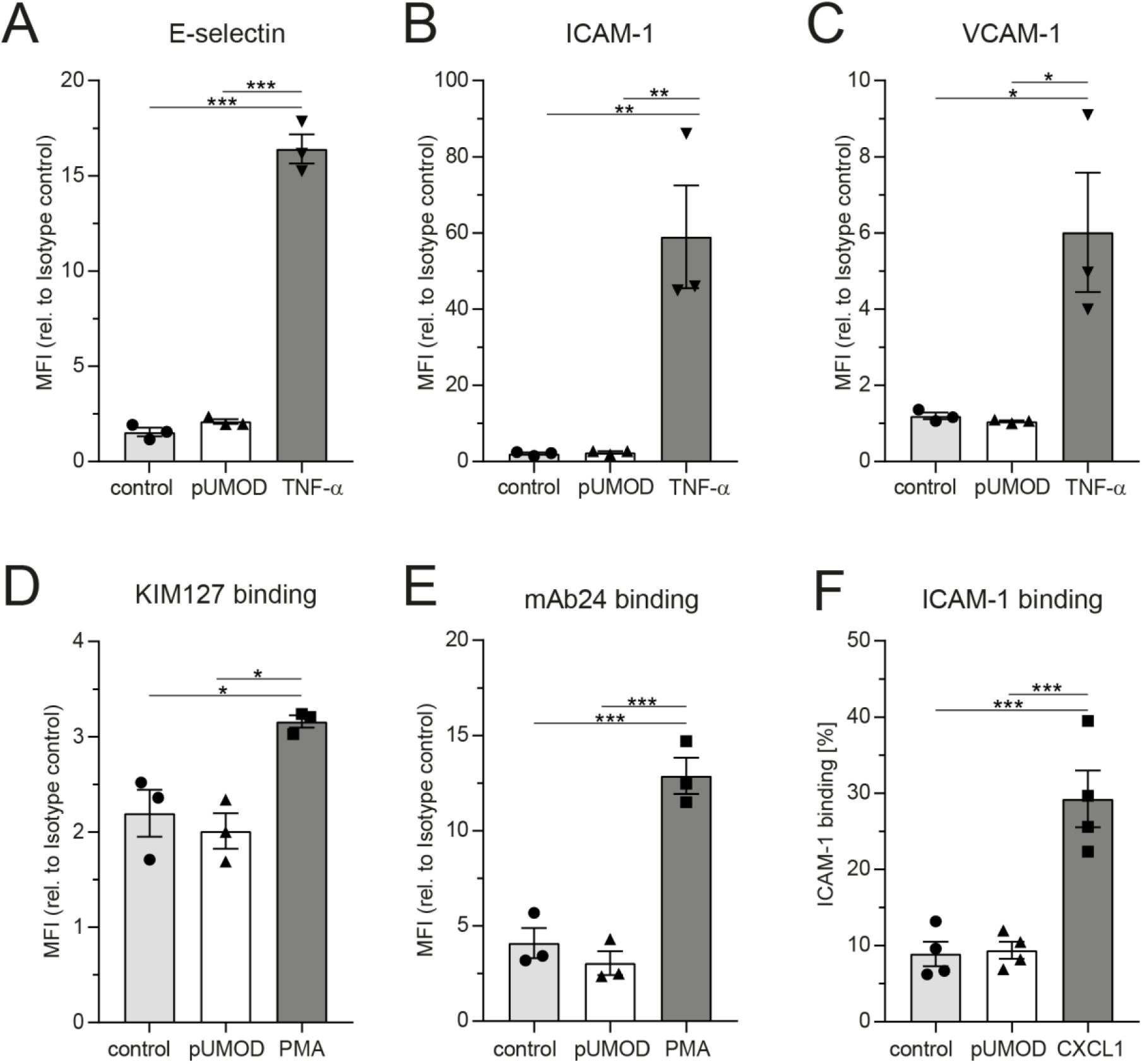
pUMOD does not directly induce adhesion molecule expression on endothelial cells or β2 integrin activation on neutrophils. HUVEC monolayers were stimulated with pUMOD, TNF-α or vehicle control and surface expression of (**A**) E-selectin, (**B**) ICAM-1 and (**C**) VCAM-1 was determined (n=3 independent experiments, one-way ANOVA, Tukey’s multiple comparison, MFI=Mean fluorescence intensity). Isolated human neutrophils were stimulated with pUMOD, PMA or vehicle control and (**D**) LFA-1 intermediate activation and (**E**) LFA-1 intermediate and fully activation state was analyzed (n=3 independent experiments, one-way ANOVA, Tukey’s multiple comparison). (**F**) Bone marrow derived murine neutrophils were stimulated with pUMOD, CXCL1 or vehicle control and binding capacity of soluble ICAM-1 to the cells was quantified (n= 4 mice per group, one-way ANOVA, Tukey’s multiple comparison). *: p≤0.05, **: p≤0.01, ***: p≤0.005, data is presented as mean±SEM.

### pUMOD does not directly induce adhesion molecule expression on endothelial cells or β2 integrin activation on neutrophils

As pUMOD application into the scrotum of WT mice induced a strong inflammatory response with functional upregulation of E-selectin on the inflamed endothelium, we aimed to investigate whether pUMOD is able to directly induce upregulation of proinflammatory adhesion molecules on primary human umbilical vein endothelial cells (HUVECs) *in vitro*. For that purpose, we stimulated a monolayer of HUVECs with pUMOD, PBS, or TNF-α for six hours. Under control conditions (PBS), HUVECs did not constitutively express VCAM-1 and only small amounts of ICAM-1 and E-selectin, as indicated by FACS analysis (**Figure 2A-C**). TNF-α stimulation in turn strongly upregulated surface levels of E-selectin, ICAM-1, and VCAM-1. Interestingly, pUMOD stimulation of HUVEC did not lead to the upregulation of non of the tested surface molecules suggesting that pUMOD does not directly affect expression levels of these rolling and adhesion related molecules on endothelial cells. This was surprising, as UMOD had been described to be a ligand for TLR4 (31). Stimulation of TLR4 through its ligand LPS is known to induce upregulation of E-selectin, ICAM-1, or VCAM-1 on the surface of HUVECs (32, 33) and β2 integrin activation on neutrophils (34). Therefore, we also analyzed potential effects of pUMOD on the activation status of β2 integrins on human and murine neutrophils. We used isolated human neutrophils from healthy donors and stimulated the cells with pUMOD, PMA, or PBS (control). The activation status of β2 integrins was analyzed by flow cytometry using the integrin activation-specific antibodies KIM127 and mAB24 (35). In contrast to PMA, pUMOD was not able to convert β2 integrins into its activated forms (**Figure 2D and 2E**). In addition, surface expression levels of Leukocyte function-associated antigen-1 (LFA-1) and total macrophage-1 antigen (Mac-1) were not affected by pUMOD (**Supplemental Figure 3**) suggesting that pUMOD does not directly stimulate β2 integrin activation. In a second set of experiments, bone marrow derived murine neutrophils were stimulated with pUMOD or recombinant mouse CXCL1 to induce GPCR mediated integrin activation. In line with the finding in human neutrophils, pUMOD stimulation did not activate β2 integrins on murine neutrophils while CXCL1 treated cells showed increased binding of recombinant murine ICAM-1 illustrating β2 integrin activation (**Figure 2F**). These *in vitro* findings indicate that pUMOD does not directly induce the expression of key adhesion molecules, neither on endothelial cells nor on neutrophils, but rather exerts its proinflammatory effects through indirect mechanisms. In addition, the experiments also exclude TLR4 as pUMOD receptor on endothelial cells and neutrophils, as previously suggested (31) in our experimental setting.

**Figure 3.**
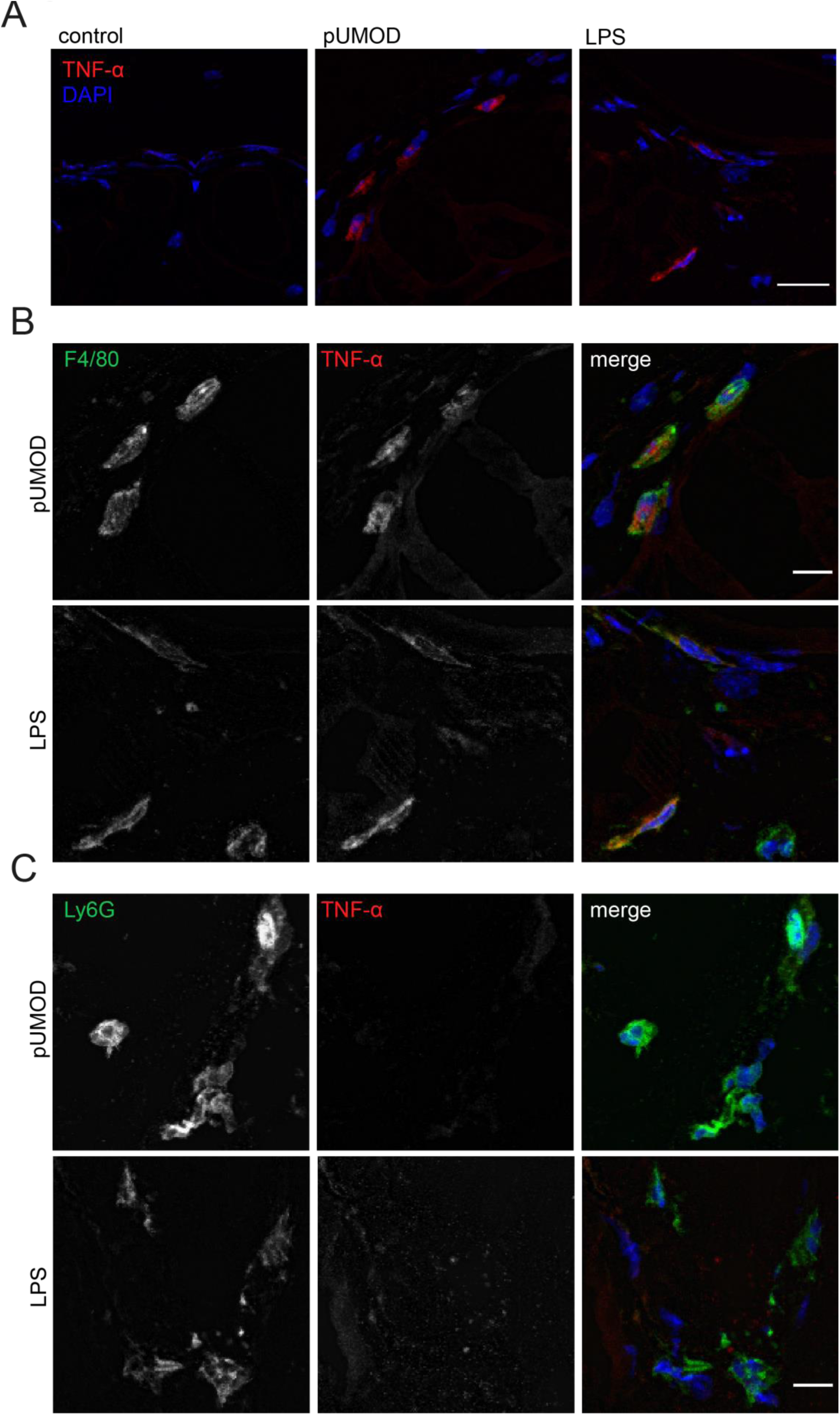
pUMOD induces TNF-α expression of F4/80 positive macrophages in the cremaster muscle *in vivo*. Cryo sections of normal saline (control), LPS and pUMOD stimulated cremaster muscles of WT mice were stained for (**A**) TNF-α expression (representative micrographs of n=3 mice per group, scale bar=20µm; red: TNF-α, blue: DAPI), (**B**) macrophages (F4/80) (representative micrographs of n=3 mice per group, scale bar=10µm, green: F4/80, red: TNF-α, blue: DAPI) and (**C**) neutrophils (Ly6G) (representative micrographs of n=3 mice per group, scale bar=10µm, green: Ly6G, red: TNF-α, blue: DAPI). Data is presented as representative images.

### pUMOD induces TNF-α expression of F4/80 positive macrophages in the cremaster muscle *in vivo*

To further explore how pUMOD might mediate its proinflammatory effects as observed in our *in vivo* cremaster muscle model, we investigated the ability of pUMOD to induce TNF-α release by tissue macrophages. An earlier report has shown that UMOD induced TNF-α secretion in human monocytes (36) and dendritic cells (31) and in bone marrow derived murine monocytes *in vitro* (31). Furthermore, intravenous administration of UMOD resulted in high serum TNF-α levels in WT mice (31). To test for pUMOD induced TNF-α release, we injected pUMOD or LPS into the scrotum of WT mice. Two hours later, we dissected and cryo-conserved cremaster muscles and stained 10µm thick frozen slices with an antibody against murine TNF-α. pUMOD as well as LPS strongly induced TNF-α expression compared to normal saline injection (control) **(Figure 3A)**. To define the cellular source of TNF-α production, we used antibodies against F4/80 and Ly6G. Application of pUMOD strongly induced TNF-α expression in F4/80 positive macrophages (**Figure 3B**), whereas it was completely absent in Ly6G expressing neutrophils (**Figure 3C**). These findings demonstrate that pUMOD stimulates F4/80 positive macrophages to produce TNF-α, thereby triggering inflammation and leukocyte recruitment.

### Intrascrotal pUMOD application potentiates the effects of TNF-α in the acute inflammation model of the mouse cremaster muscle *in vivo*

To elucidate whether pUMOD affects leukocyte recruitment exclusively by activating TNF-α production in F4/80 positive macrophages or whether it may have additional, TNF-α independent effects on leukocyte recruitment, we injected recombinant murine TNF-α alone or in combination with pUMOD into the scrotum of WT mice and analyzed leukocyte recruitment two hours after onset of inflammation. pUMOD did not alter TNF-α induced leukocyte rolling, leukocyte adhesion and leukocyte rolling velocities in postcapillary venules of the mouse cremaster (**Figure 4A-C**). Surprisingly, pUMOD potentiated the effect of TNF-α on leukocyte extravasation. The number of extravasated neutrophils as analyzed in cremaster muscle whole mounts was significantly increased when pUMOD was applied concomitantly with TNF-α compared to TNF-α application alone (**Figure 4D and 4E**). Of note, hemodynamic parameters did not differ between the groups (**Supplemental Table 3**). These experiments suggest a role for pUMOD in the promotion of neutrophil extravasation *in vivo* independent of the effects triggered by TNF-α.

**Figure 4.**
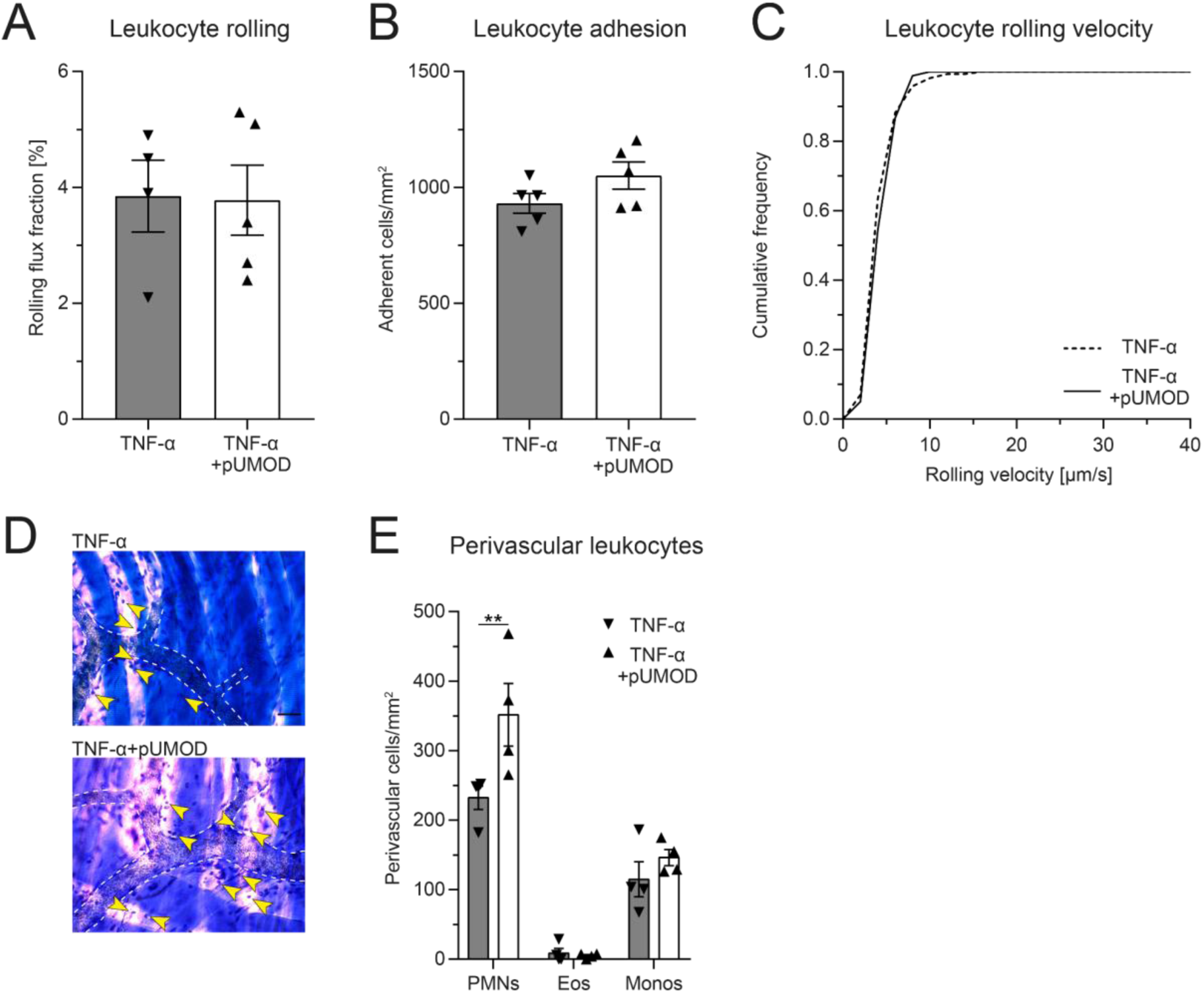
Intrascrotal pUMOD application potentiates the effects of TNF-α in the acute inflammation model of the mouse cremaster muscle *in vivo*. WT mice were injected i.s. with TNF-α or a combination of TNF-α and pUMOD. Two hours after stimulation, intravital microscopy of postcapillary venules of the cremaster muscle was performed and (**A**) rolling flux fraction and (**B**) number of adherent leukocytes was analyzed in 34 (TNF-α) and 35 (TNF-α+pUMOD) venules of n=5 mice per group. (**C**) Leukocyte rolling velocity was determined in n=170 (TNF-α) and 180 (TNF-α+pUMOD) cells of 5 mice per group. Concomitant application of TNF-α and pUMOD did not modify rolling flux fraction, leukocyte adhesion or rolling velocities compared to TNF-α administration (unpaired students t-test). (**D**) Cremaster muscles were stained with Giemsa (representative micrographs of n=4 mice per group. Arrow bars indicate perivascular leukocytes, (scale bar=30µm). (**E**) Number of perivascular neutrophils (PMNs), eosinophils (Eos) and monocytes (Monos) was analyzed in 28 perivascular regions of n=4 mice per group. Concomitant application of TNF-α and pUMOD increased the number of perivascular neutrophils compared to TNF-α administration (two-way repeated measurements ANOVA, Sidak’s multiple comparison). **: p≤0.01, data is presented as mean±SEM, cumulative frequency or representative images.

### pUMOD facilitates neutrophil transmigration across a HUVEC monolayer and increases vascular permeability *in vitro* and *in vivo*

To elucidate a potential direct influence of pUMOD on neutrophil transmigration, which is independent of its effect on F4/80 positive macrophages in more detail, we allowed HUVECs to grow to confluence on transwell filters and stimulated the monolayer with pUMOD or PBS (control) for five hours. Afterwards we allowed isolated human neutrophils to transmigrate through the monolayers in absence (w/o) or presence of CXCL8 in the lower compartment of the transwell system. As expected, CXCL8 induced efficient neutrophil transmigration compared to w/o control (**Figure 5A**). Interestingly, pre-stimulation of the monolayer with pUMOD facilitated neutrophil transmigration with or without CXCL8, demonstrating a direct effect of pUMOD on transendothelial migration of neutrophils and therefore on neutrophil diapedesis and extravasation. Next, we tested the effect of pUMOD on vascular permeability of endothelial cells by measuring the electrical impedance over a HUVEC monolayer. Both, pUMOD and TNF-α significantly decreased electrical impedance of HUVEC monolayers compared to control (**Figure 5B**). Interestingly, concomitant application of pUMOD and TNF-α did not further reduce electrical impedance of the monolayer. These results demonstrate that pUMOD increases vascular permeability of endothelial cells similarly to TNF-α. Finally, we wanted to confirm pUMOD-induced changes in vascular permeability *in vivo*. For this approach, we injected pUMOD into the scrotum of WT mice. Two hours later, we prepared the mouse cremaster muscle for intravital imaging, injected FITC-dextran through the carotid artery catheter and observed changes in fluorescence intensity in the tissue surrounding cremaster muscle postcapillary venules over time. In line with the *in vitro* findings, i.s. injection of pUMOD resulted in a significant increase of vascular permeability *in vivo* compared to injection of normal saline (control) (**Figure 5C and 5D**). Taken together, these results clearly show a direct proinflammatory function of pUMOD on the vasculature by increasing vascular permeability and promoting neutrophil transmigration *in vitro* and *in vivo*.

**Figure 5.**
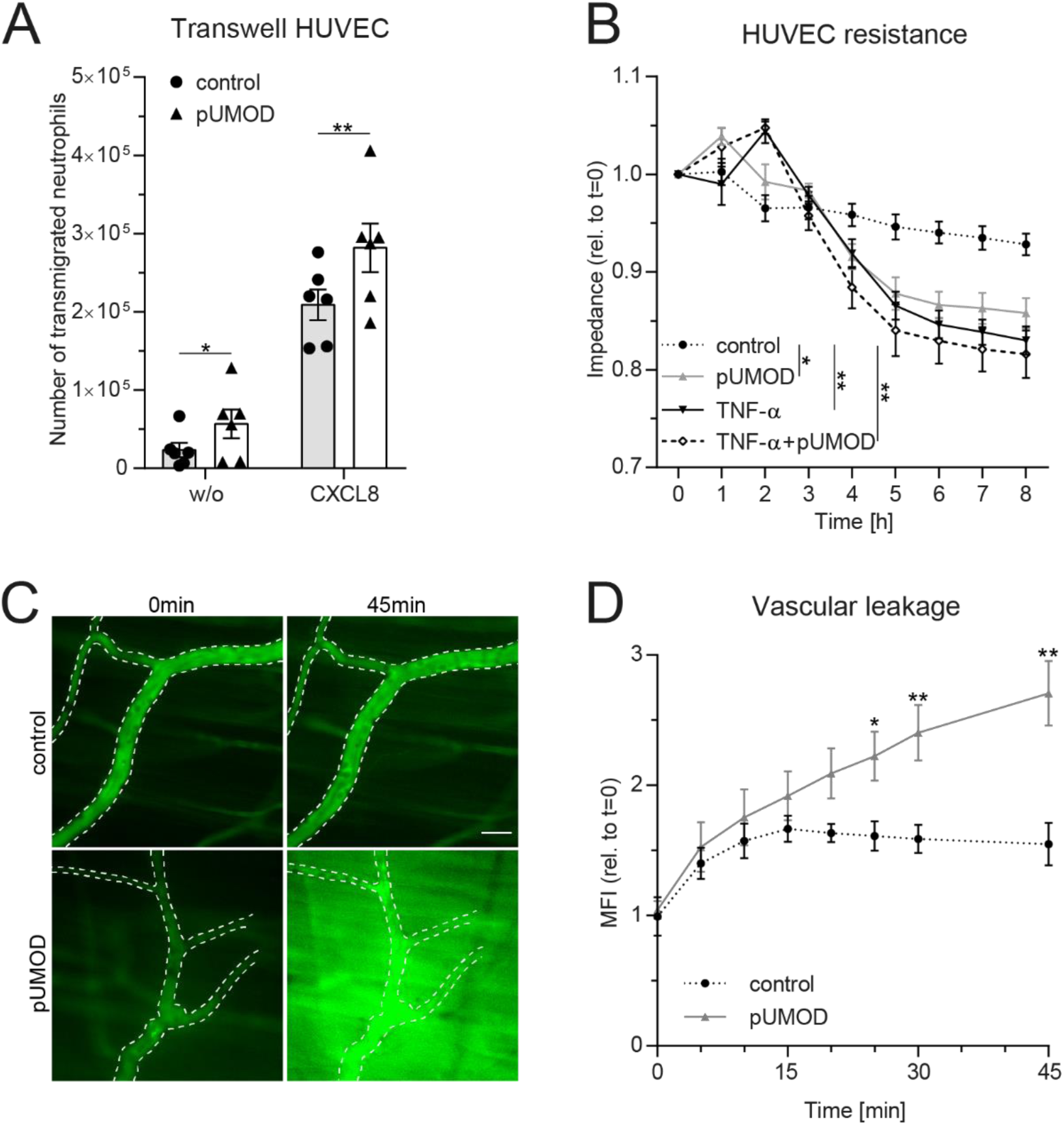
pUMOD facilitates neutrophil transmigration across a HUVEC monolayer and increases vascular permeability *in vitro* and *in vivo*. (**A**) Number of human neutrophils transmigrating along a CXCL8 or HBSS (w/o) gradient through a HUVEC monolayer pre-stimulated with pUMOD or vehicle control (control). (**B**) Electrical impedance of HUVEC monolayers were measured over time in the presence of pUMOD, TNF-α, a combination of pUMOD+TNF-α or vehicle control (control) (n=5 independent experiments, repeated one-way ANOVAs, Tukey’s multiple comparisons). (**C**) Intravital microscopy of WT mice pretreated i.s. with pUMOD or normal saline (control). Through systemic injection of FITC-dextran, vascular leakage was assessed over time (representative micrographs of n=6 mice per group, scale bar=30µm) and (**D**) changes in mean fluorescence intensity (MFI) were quantified (n=6 mice per group, unpaired student’s t-test). *: p≤0.05, **: p≤0.01, data is presented as mean±SEM.

### Neonatal obstructive nephropathy induces intra- and extratubular UMOD accumulation accompanied by tubular atrophy and leukocyte infiltration

As UMOD accumulation within renal tubules has been reported during normal neonatal development, we investigated UMOD expression in neonatal kidneys after unilateral ureteral obstruction (UUO) thereby linking a potential accumulation of UMOD at extratubular sites to leukocyte infiltration, which is commonly observed in neonatal UUO (37, 38). First, we performed Western Blot analysis in sham (control) and UUO operated neonatal kidneys. Expression levels of UMOD significantly increased after UUO compared to sham operated controls (**Figure 6A and 6B)**. Immunohistochemical analysis displayed tubular dilatation and UMOD accumulation at intra- and extratubular sites of the dilated tubule, while controls displayed normal UMOD distribution exclusively insight the epithelium and the tubular lumen **(Figure 6C)**. In addition, UUO led to a significant increase of tubular atrophy in proximal and distal tubules (**Figure 6D and 6E**) going along with infiltration of CD45^+^/CD11b^+^ leukocytes (myeloid cells) as shown by FACS analysis (**Figure 6F**).

**Figure 6.**
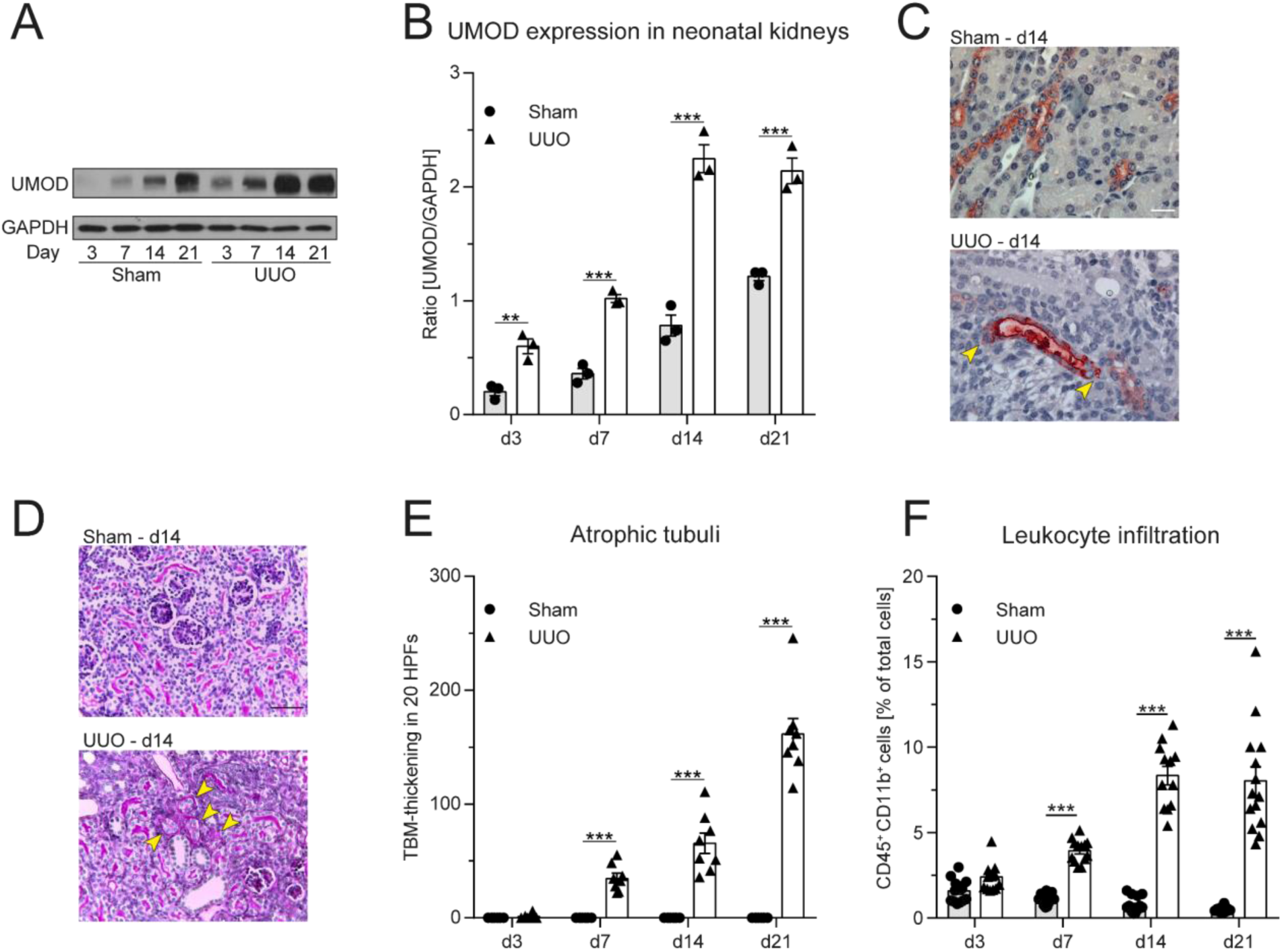
Neonatal obstructive nephropathy induces intra- and extratubular UMOD accumulation accompanied by tubular atrophy and leukocyte infiltration. (**A**) Changes in total UMOD protein levels over time in sham and UUO operated neonatal kidneys were analyzed (representative blot) and (**B**) quantified (n=3 independent experiments, two-way repeated measurements ANOVA, Sidak’s multiple comparison). (**C**) Intra- and extratubular (yellow arrows) UMOD accumulation (scale bar=20µm) and (**D**) PAS staining in sham and UUO operated neonatal kidneys was analyzed (representative micrographs, scale bar=200µm; arrow bars indicate atrophic tubuli). (**E**) Tubular atrophy (n=8 mice per group, two-way repeated measurements ANOVA, Sidak’s multiple comparison; TBM: tubular basement membrane; HPF: high-power field) and (**F**) myeloid cell infiltration was assessed in sham and UUO operated neonatal kidneys (n=12-16 mice per group, two-way ANOVA, Sidak’s multiple comparison). **: p≤0.01, ***: p≤0.005, data is presented as mean±SEM or representative images.

### UMOD casts in patients suffering from monoclonal gammopathy with renal significance, IgA nephropathy and interstitial nephritis co-localize with immune cell infiltrates

Finally, we analyzed UMOD expression and localization in patients suffering from monoclonal gammopathy with renal significance, IgA nephropathy and interstitial nephritis. For this approach, we used human renal biopsies and stained tissue sections with PAS, CAE and a specific anti human UMOD antibody **(Figure 7)**. We detected UMOD casts within the kidney, surrounded by infiltrates of neutrophils indicating a proinflammatory function of pUMOD if located outside the tubular lumen.

**Figure 7.**
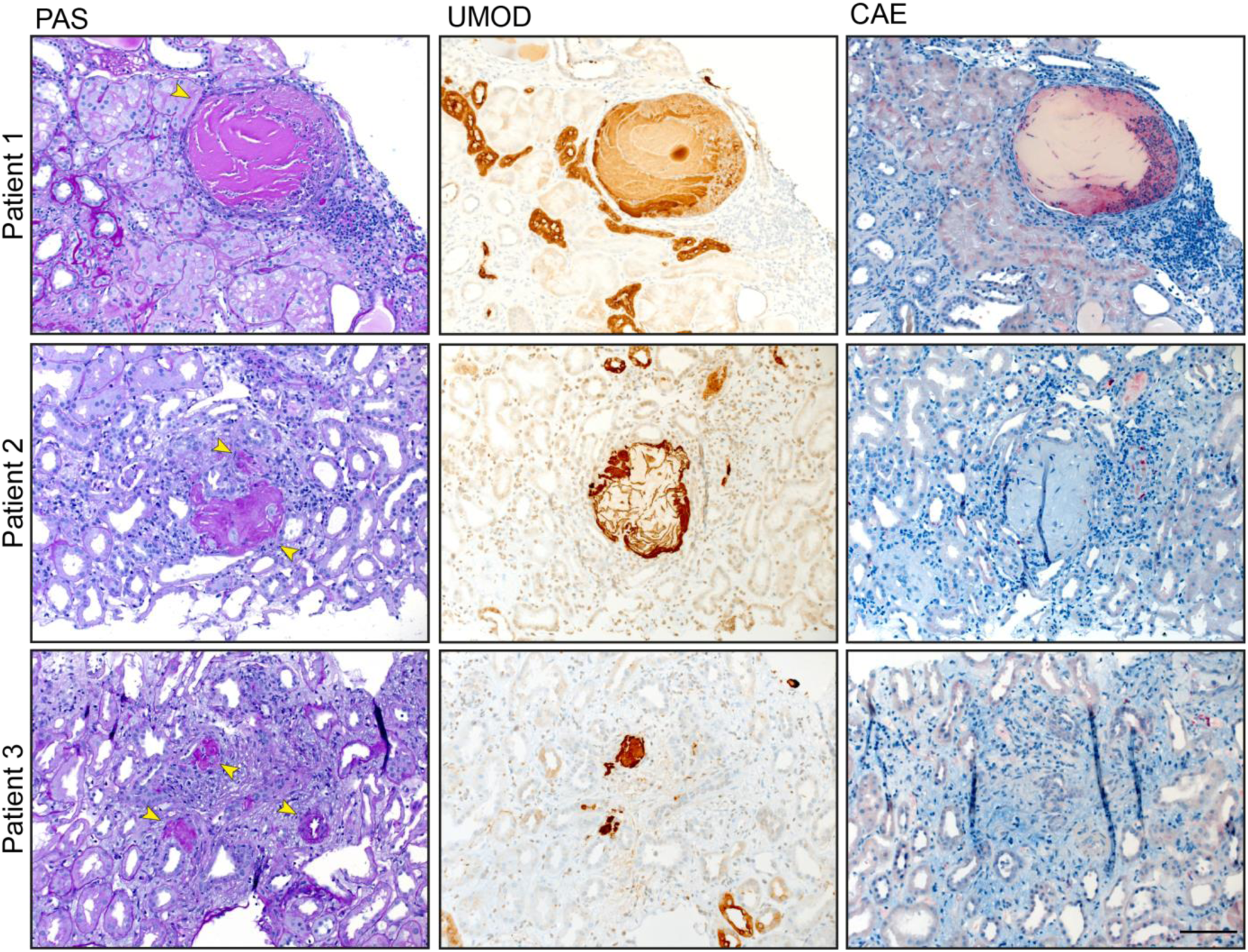
UMOD casts colocalize with neutrophil infiltrates. We detected UMOD casts in the kidneys of three patients suffering from monoclonal gammopathy with renal significance, IgA nephropathy, and interstitial nephritis with arterial hypertension, respectively. PAS and antibody staining display UMOD localization (arrows). Neutrophil infiltrates, stained with anti CEA antibody, were found in close proximity to UMOD casts (scale bar=100µm)

## DISCUSSION

UMOD is a pleiotropic protein secreted by tubular epithelial cells of the thick ascending limb of the loop of Henle and to a lesser degree the early distal convoluted tubule. Here, we investigated a potential proinflammatory role of extratubular UMOD using pUMOD isolated from human urine. Stimulation of extrarenal tissue (mouse cremaster muscle and peritoneum as *in vivo* models of acute microvascular inflammation) with pUMOD *in vivo* led to a strong inflammatory response reflected by marked recruitment of neutrophils and inflammatory monocytes. Complemented with various additional *in vivo* and *in vitro* assays we demonstrate that pUMOD directly modulates the endothelial permeability and stimulates the generation of TNF-α from tissue macrophages. This in turn triggers the activation of vascular endothelial cells leading to the upregulation of adhesion molecules including E-selectin and the subsequent influx of neutrophils and inflammatory monocytes into the inflamed tissue.

Neutrophils and inflammatory monocytes belong to the innate immune system and are key players in the first line of defense against invading pathogens or during tissue damage. They exit the vasculature into inflamed tissue through a series of consecutive adhesion and activation steps (39). Several *in vitro* studies have predicted a role of UMOD on innate immune cell functions. UMOD was described to induce TNF-α secretion and tissue factor expression in human monocytes (36), to trigger dendritic cell maturation via TLR4 and NF-κB activation (31) and to activate respiratory burst, release of proteinases, degranulation and phagocytosis in neutrophils (40-42). Additionally, UMOD was reported to enhance CXCL8 expression, to dampen CD62L expression in human granulocytes (43) and to stimulate the NLRP3 inflammasome in human monocytes, resulting in IL-1β release and cellular death (44). UMOD had also been reported to facilitate neutrophil transepithelial migration *in vitro* by using a stably transfected UMOD expressing epithelial cell line (45). Furthermore, a proinflammatory role of UMOD had also been described through the observation that intravenous administration of UMOD results in systemic inflammation (31). On the other hand, UMOD has been reported to limit inflammation by suppressing the production of proinflammatory cytokines and chemokines (46-48). Micanovic et al. found a negative regulatory role of UMOD on granulopoiesis and systemic neutrophil homeostasis (49). Interestingly, the same group described that proliferation and phagocytic activity of mononuclear phagocytes can be enhanced by UMOD while at the same time, loss of UMOD leads to an aggravated immune response in a renal ischemia-reperfusion injury model (14). Although these findings on the role of UMOD during inflammation seem at least in part conflicting, Micanovic and colleagues provide an attractive explanation by proposing that UMOD functions can be assigned to two different forms of UMOD described *in vivo*, namely the monomeric and the polymeric form. Under homeostatic conditions, these two UMOD forms localize to two different body compartments after being released from tubular epithelial cells (2, 14, 30). The polymeric form is the predominant form found in the tubular lumen and urine, where it regulates water and electrolyte balance and protects the kidney from bacterial infections and kidney stone formation (2). Additionally, tubular epithelial cells release, although to a lesser extent, UMOD to the basolateral site into the renal parenchyma. Basolaterally released UMOD does not polymerize, but instead is transported into the bloodstream where it circulates primarily in its monomeric form and exerts its systemic homeostatic functions (14, 30). Therefore, under physiological conditions, the polymeric form is exclusively found in the urinary tract and is not exposed to tissues outside of the tubular lumen. Any contact of pUMOD with renal tissue outside the lumen only occurs under pathological conditions, i.e. in case the epithelial barrier with its tubular basement membrane is disrupted. Pathophysiological deposition of UMOD, so called cast nephropathy, is known for a long time as a common renal complication in the pathology of multiple myeloma, where free light chains present in the ultra-filtrate bind to UMOD and co-precipitate to form casts (50). These casts are supposed to induce tubular atrophy, progressive interstitial inflammation and fibrosis, resulting in renal complications, like acute kidney injury (25). We detected UMOD casts beyond their appearance in multiple myeloma, in patients suffering from monoclonal gammopathy of renal significance, IgA nephropathy and interstitial nephritis, where infiltrates of neutrophils surround UMOD casts in the kidney.

For this reason, we tested whether pUMOD functions as a DAMP-like molecule triggering inflammation if reaching renal tissue outside of the urinary tract. We show that extratubular pUMOD increases vascular permeability of endothelial cells, facilitates leukocyte extravasation and activates F4/80 positive macrophages to produce proinflammatory factors. However, it is currently unknown how pUMOD activates endothelial cells and innate immune cells, thereby mounting an immune response. Several UMOD receptors have been described including TLR4, Transient receptor potential melastatin 2 (TRPM2) and TRPM6 channels (13, 16, 31). TRPM6 is neither expressed on endothelial cells nor on neutrophils, suggesting TLR4 and TRPM2 as putative pUMOD interaction partners responsible for the described proinflammatory effects. TLR4 ligation with LPS is known to upregulate E-selectin and ICAM-1 expression on HUVEC (32, 33). Using pUMOD as putative ligand for TLR4 on HUVEC, we were not able to induce upregulation of E-selectin, indicating that endothelial TLR4 is not a target of pUMOD in our experimental setting. Further support that pUMOD does not functionally engage TLR4 can be deduced from our *in vitro* experiments on isolated neutrophils where we showed that pUMOD stimulation of neutrophils did not activate β2 integrins on the neutrophil surface. In contrast, recent work from our group demonstrated that the TLR4 ligands LPS and S100A8/A9 induce activation of β2 integrins in neutrophils (34). TRPM2, the other reported pUMOD receptor was shown to regulate endothelial barrier function (51). In addition, TRPM2 is expressed on macrophages(52) and neutrophils (53). Therefore, it is tempting to speculate that TRPM2 may be involved in pUMOD-induced changes in endothelial barrier function, macrophage activation and neutrophil/monocyte recruitment as described here. Nevertheless, the precise underlining mechanism of how pUMOD modulates endothelial and innate immune cells on a molecular level and which receptor/receptors are involved is still unclear and needs further investigation.

UMOD is highly expressed in the developing kidney and urinary tract obstruction during kidney development leads to leukocyte recruitment, inflammation and interstitial fibrosis in the neonatal kidney (37, 54). Therefore, we investigated UMOD expression, localization and immune cell infiltration in a neonatal model of unilateral ureteral obstruction (UUO). We show that UUO increases UMOD expression that localizes within and outside the tubular lumen. UUO causes tubular cell death and tubular atrophy leading to changes in tubular basement membrane integrity, which might favor the leakage of pUMOD into extratubular renal parenchyma. We postulate that in neonatal UUO pUMOD reaches extratubular renal tissue and induces recruitment of innate immune cells and interstitial inflammation in the neonatal kidney with obstruction. Our findings are in line with observations showing that UMOD protein levels increased in kidneys of adult mice following UUO and uromodulin-deficiency reduced inflammation and kidney injury in this model (26). Interestingly, interstitial UMOD deposition could not be detected in substantial quantities within *Umod* ^+/+^ mice with UUO in their study. However, this does not exclude a proinflammatory role of extratubular pUMOD in this process. In fact, low quantities of extratubular pUMOD undetectable by immunohistology may be sufficient (and of critical importance) to activate endothelial cells and tissue resident macrophages in order to mount an inflammatory response in case of lost urinary tract barrier integrity.

Taken together, we identified a strong proinflammatory function of extratubular pUMOD with the potential to induce TNF-α secretion from kidney-resident macrophages and to modify endothelial cell permeability, thereby promoting innate immune cell (myeloid cell) recruitment and enhancing inflammation. This might be a scenario during acute and chronic renal diseases where luminal pUMOD accumulation and rupture of the tubular basement membrane leads to pUMOD release into the extratubular renal parenchyma. Extratubular pUMOD might then provide an important local DAMP-like signal, which sets in motion an inflammatory response to repair damaged tissue and restore tissue homeostasis.

## METHODS

### Animals

C57BL/6 wildtype (WT) mice were obtained from Janvier Labs (Saint Berthevin) or from Charles River Laboratories (Sulzfeld). All mice were maintained at the Walter Brendel Centre of Experimental Medicine, LMU, Munich, or at the core facility animal models at the Biomedical Center, LMU, Planegg-Martinsried, Germany and at the zentrale Versuchstierhaltung Innenstadt (ZVH), LMU Munich, Germany. 8-25 weeks old male and female mice were used for all experiments. Neonatal C57BL/6 wildtype (WT) mice were used for unilateral ureteral obstruction experiments.

### Antibodies

To block E-selectin dependent leukocyte rolling *in vivo*, mice were injected with rat anti mouse E-selectin antibody (clone 9A9, In Vivo, 30µg/mouse). Following antibodies were used for UMOD Western blot: sheep anti mouse UMOD (R&D Systems, 1:1000), rabbit anti sheep IgG (Southern Biotech, 1:2000), for UMOD immunohistochemistry in UUO model: rat anti UMOD (R&D Systems, 1:100) and for UMOD immunohistochemistry of human patient samples: mouse anti human UMOD (kindly provided by Jürgen Scherberich(55), 1:800). CAE staining was used to determine granulocytes. Following antibodies were used for flow cytometry (all 5µg/ml, except indicated differently): rat anti mouse CD45 (clone 30-F11, BioLegend), rat anti mouse CD11b (clone M1/70), rat anti mouse CD115 (clone AFS98, BioLegend), rat anti mouse Gr1 (clone RB6-8C5, BioLegend), mouse anti human CD66b (clone G10F5, BioLegend) and CD66abce (clone TET2, Miltenyi Biotec), mouse anti human CD15 (clone W6D3, BioLegend or clone VIMC6, Miltenyi Biotec), mouse anti human CD62E (clone HAE-1f, BioLegend), mouse anti human IgG_1_ isotype control (MOPC-21, BioLegend), mouse anti human CD54 (clone HA58, BioLegend), mouse anti human IgG_1_ isotype control (clone 11711, R&D Systems), mouse anti human CD106 (clone 4B2, R&D Systems), secondary goat anti mouse-PE (Pharmingen),mouse anti human CD18 high affinity confirmation (clone 24, Hycult Biotech, 10μg/ml), mouse anti human CD18 extended confirmation (clone KIM127, Invivo, 10µg/ml), mouse anti human CD11a (clone HI111, BioLegend, 10μg/ml), mouse anti human CD11b (clone ICRF44, BioLegend, 10μg/ml), mouse anti human IgG_1_ isotype control (clone 11711, R&D Systems, 10μg/ml), rat anti mouse Ly6G (clone 1A8, BioLegend), goat anti human Fcγ–biotin (eBioscience) and streptavidin–PerCP-Cy5.5 (BioLegend). For immunofluorescence, we used the following antibodies: rat anti mouse F4/80 (Invitrogen, 1:200), rabbit anti TNF-α (abcam, 1:400), and rat anti mouse Ly6G (clone 1A8, BioLegend, 1:100). As secondary antibodies we used: donkey anti rat Alexa 488 (Invitrogen, 1:400) and goat anti rat Alexa 546 (Invitrogen, 1:400). DAPI (4,6-diamidino-2-phenylindole) was used as a nuclear counterstain.

### Transmission electron microscopy

Transmission electron microscopy was performed as described previously (44). All samples were stored at 4 degrees until further processing. In brief, a 30µl drop of human or mouse uromodulin protein in dH20 was placed on a formvar-coated copper grid (Science Services) and fixed with 2,5% Glutaraldehyde (Science Services) for 1 minute, washed in dH20 for 1 minute and stained with 0,5% UranylLess (Science Services) in dH20 for 1 minute and air dried. Imaging was carried out using the JEOL -1200 EXII transmission electron microscope (JEOL, Akishima) at 60 kV. Images were taken using a digital camera (KeenViewII; Olympus) and processed with the iTEM software package (anlySISFive; Olympus).

### Animal model of acute inflammation in the mouse cremaster muscle

Intravital microscopy of the mouse cremaster muscle was performed as previously described (34). Briefly, WT mice received an intrascrotal (i.s.) injection of either polymerized UMOD (pUMOD, BBI Solutions), human serum albumin (hALB, Sigma-Aldrich), murine recombinant polymerized UMOD (murine pUMOD, LSBio), murine serum albumin (murine ALB, Sigma-Aldrich) (all 10µg/mouse in 200µl 0.9%NaCl) or vehicle (control; 0.9% NaCl). The concentration used for pUMOD were chosen in accordance to previous studies (31). We also tested for potential LPS contamination of pUMOD and found no indication of TLR4 activation in functional assays (Figure 2A-F). In a second set of experiments, WT mice received an i.s. injection of recombinant murine (rm)TNF-α (R&D Systems, 500ng/mouse) or a concomitant injection of rmTNF-α and pUMOD (10µg/mouse). Two hours after i.s. injection, mice were anesthetized and the carotid artery was cannulated for determination of the white blood cell count (ProCyte Dx; IDEXX Laboratories) and/or for the application of E-selectin-blocking antibodies (30µg/mouse). The cremaster muscle was dissected and intravital microscopy was carried out on an OlympusBX51 WI microscope, equipped with a 40x objective (Olympus, 0.8NA, water immersion objective) and a CCD camera (Kappa CF 8 HS). During the experiment, the muscle was constantly superfused with thermo-controlled bicarbonate buffer and postcapillary venules were recorded using VirtualDub software for later off-line analysis. Centerline velocity of each venule was measured with a dual photodiode (Circusoft Instrumentation). Rolling flux fraction (number of rolling cells normalized to complete leukocyte flux (28)), number of adherent cells/mm^2^, leukocyte rolling velocities, vessel diameter and vessel length was determined on the basis of the generated movies using Fiji software (56). In another set of experiments, cremaster muscles were removed, fixed with paraformaldehyde (4% w/v (Applichem) in phosphate-buffered solution (PBS)) and used for Giemsa-staining (Merck) to calculate the number of perivascular neutrophils, eosinophils and monocytes. The analysis of perivascular leukocytes was carried out at the core facility BioImaging of the Biomedical Center with a Leica DM2500 microscope, equipped with a 100x objective (Leica, 1.4NA, oil immersion) and a Leica DMC2900 CMOS camera. In an additional set of experiments, cremaster muscles were removed, frozen in OCT embedding medium (Sakura Finetec) and stored at −80°C until further processing. Embedded frozen tissues were sectioned at 10μm with a Leica cryostat and fixed with ice-cold acetone for 10 minutes. Sections were used for an antigen-specific antibody staining procedure. For *in vivo* analysis of endothelial permeability, mice received an i.s. injection of pUMOD (10µg/mouse) or vehicle (control; 0.9% NaCl). Two hours later, mice were anesthetized and the carotid artery was cannulated for the application of fluorescein isothiocyanate (FITC)–dextran (Sigma-Aldrich, 150kD) as described elsewhere (57). Briefly, 3 postcapillary venules and the surrounding tissue of the dissected mouse cremaster muscle were recorded for 45 minutes with the help of an Axio Scope.A1 microscope, equipped with a 488nm LED light source (Colibiri 2) a long pass emission filter (LP 515), a 20x water immersion objective 0.5NA and a AxioCam Hsm digital camera (Zeiss MicroImaging). Mean fluorescence intensities of 6 randomly chosen regions of interest (ROI, 50×50µm^2^) were analyzed using ImageJ. Each ROI was at least 50µm in distance from the respective vessel.

### Animal model of acute inflammation in the mouse peritoneum

WT mice received an intraperitoneal (i.p.) injection of either pUMOD (10µg/mouse in 800µl 0.9%NaCl) or vehicle (control; 0.9% NaCl). 6h later, peritoneal lavage was performed using 7ml ice-cold PBS. Cells were collected, stained and number of extravasated neutrophils and inflammatory monocytes were counted by flow cytometry (CytoFLEX, Beckman Coulter) using Flow-Count Fluorospheres (Beckman Coulter) and analyzed with FlowJo Analysis Software. Neutrophils and inflammatory monocytes were defined as CD45^+^/CD11b^+^/Gr1^+^/CD115^-^and CD45^+^/CD11b^+^/Gr1^+^/CD115^+^ cells, respectively.

### Immunofluorescence of the mouse cremaster muscle

Immunofluorescence was carried out using 5µm thick cryosections of mouse cremaster muscles. Briefly, cryosections were washed thoroughly in PBS before and after each incubation step. Nonspecific binding sites were blocked by incubation in 20% normal goat serum (NGS) and 5% bovine serum albumin/PBS (BSA, Fraction V; both Sigma-Aldrich). Cryosections were stained with anti TNF-α, anti F4/80 and anti Ly6G antibodies, followed by secondary antibody staining. Sections were counterstained with DAPI and evaluated using confocal microscopy at the core facility bioimaging of the Biomedical Center with an inverted Leica SP8 microscope.

### Isolation and cell culture of HUVECs

HUVECs were purchased from Promocell (HUVEC cryopreserved, pooled) or freshly isolated from the umbilical cord vein by collagenase A digestion (1mg/ml; Roche Diagnostics), as previously described (58). Cells were grown in Endothelial cell Growth Medium supplemented with SupplementMix (Promocell) and DMEM supplemented with 20% FCS and 1% penicillin/streptomycin (PAA Laboratories) in a 1:1 ratio in standard cell culture dishes. At confluence, cells were harvested using trypsin/EDTA (PAA Laboratories). Cells from passage II or III were used for further experiments.

### Transwell Migration Assay

HUVECs were cultured on Transwell inserts (Falcon, 8μm pore size) to confluence. Confluent monolayers were stimulated with pUMOD (3µg/ml) or PBS (control) for 5h before being washed. Human neutrophils were isolated from healthy volunteer blood donors using Polymorphprep (AXI-SHIELD) as previously described (34). Isolated neutrophils were applied in the upper compartment of the Transwell system onto the HUVEC monolayer and allowed to migrate for 1h at 37°C in HBSS puffer containing 0.1% glucose, 1mM CaCl_2_, 1mM MgCl_2_, 0.25% BSA and 10mM Hepes (Sigma-Aldrich), pH7.4. Recombinant CXCL8 (Peprotech, 100ng/ml) was used as chemoattractant, HBSS puffer alone was used as negative control (w/o). Numbers of transmigrated neutrophils was counted by flow cytometry (Gallios, Beckman Coulter) using Flow-Count Fluorospheres (Beckman Coulter) and analyzed with Kaluza Flow Analysis Software (Beckman Coulter). Neutrophils were defined as CD66b and CD15 double positive cells.

### Surface expression levels of E-selectin, ICAM-1, and VCAM-1 on HUVEC

HUVECs (passage II or III) were cultured until confluence. Confluent monolayers were stimulated with pUMOD (3µg/ml), recombinant human TNF-α (rhTNF-α, Peprotech, 10ng/ml) or PBS (control) for 6h. Cells were washed, harvested and stained with anti E-selectin (CD62E), mouse anti human ICAM-1 (CD54) and mouse anti human VCAM-1 (CD106) antibodies and fixed with FACS lysing solution (BD Bioscience). Surface levels were determined by flow cytometry (Gallios, Beckman Coulter) and analyzed using Kaluza software (Beckman Coulter).

### mAb24 binding assay

Human neutrophils were isolated from healthy volunteer blood donors using Polymorphprep and incubated with pUMOD (3µg/ml), PMA (Sigma-Aldrich, 1nM) or HBSS buffer (unstim.) in the presence of the anti-human β2 integrin activation antibody mAB24 or isotype control at 37°C for 5 minutes. Stimulation was stopped by adding ice-cold FACS lysing solution (BD Bioscience). After fixation, cells were stained with secondary goat anti mouse antibody. Activation status of β2 integrins was determined by flow cytometry, human neutrophils were defined as CD66abce and CD15 double positive cells. In addition, total Mac-1 and total LFA-1 surface protein levels were investigated. Therefore, activated neutrophils were stained with mouse anti-human CD11b or CD11a, followed by secondary antibody.

### ICAM-1 binding assay

Soluble ICAM-1 binding assay was performed as described previously (59). Briefly, bone marrow–derived neutrophils from WT mice were enriched using a Percoll gradient (Sigma-Aldrich). Enriched neutrophils were stimulated with rmCXCL1 (100ng/ml, PeproTech), pUMOD (3µg/ml) or HBSS buffer (unstim.), in the presence of rmICAM-1 (hFc chimera, 20μg/ml, R&D Systems, pre-complexed with goat anti human Fcγ–biotin and streptavidin–PerCP-Cy5.5) for 3 minutes at 37°C in HBSS buffer. Reaction was stopped by adding ice-cold FACS lysing solution. Cells were stained with rat anti mouse Ly6G antibody and the amount of bound soluble ICAM-1 was determined by flow cytometry.

### Trans endothelial electrical resistance measurements (TEER)

8 Well-Slides (8W10E, Applied BioPhysics) were coated with collagen type 1 (rat tail, Ibidi) and HUVEC were seeded onto the slides. The slides were installed in the TEER measuring device (ECISz, Applied BioPhysics), which was positioned in an incubator with 5% CO_2_ at 37°C. HUVECs (passage II) were cultured to confluence and stimulated with 3µg/ml pUMOD and/or 10ng/ml rhTNF-α at time point 0. PBS was used as negative control. Measurements were acquired continuously at 4000Hz (ECIS Software v1.2.156.0 PC, Applied BioPhysics) for 8h. Means of two technical replicates were calculated.

### Animal model of unilateral ureteral obstruction

Newborn WT mice were subjected to complete left ureteral obstruction (UUO) or sham operation under general anesthesia with isoflurane and oxygen at the second day of life as described before (37, 54, 60). After recovery, neonatal mice were returned to their mothers until sacrifice at day 3, 7, 14, or 21 of life. Kidneys were removed and either fixed in 4% PFA for immunostaining, directly used for FACS analysis or frozen in liquid nitrogen and stored at -80°C for further experiments.

### Identification of UMOD

UMOD localization in the neonatal kidney was examined by immunohistochemistry. Paraformaldehyde-fixed, paraffin-embedded kidney sections were subjected to antigen retrieval and incubated with an anti UMOD antibody, followed by a biotinylated goat anti rat IgG antibody. Specificity was assessed through simultaneous staining of control sections with an unspecific, species-controlled primary antibody. Sections were incubated with ABC reagent, detected with DAB (Vectastain, Vector Laboratories) and counterstained with methylene blue.

### PAS staining of sham and UUO operated neonatal kidneys

Kidney sections were stained with periodic acid Schiff (PAS) to investigate tubular atrophy in UUO and sham-operated mice as described previously (37, 54, 60). Alterations of the tubular basement membrane (TBM) were determined in 20 sequentially selected fields at x400 magnification.

### FACS staining of sham and UUO operated neonatal kidneys

Preparation of renal single-cell suspensions was performed as described previously (61). The obtained cell suspension contained infiltrating leukocytes and intrinsic renal cells, the largest proportion being tubular epithelial cells. Infiltrating renal myeloid cells were quantified by flow cytometry (FACSCalibur) and analyzed as Cellquest software (BD Biosciences). Myeloid cells were defined as CD45^+^/CD11b^+^ cells and expressed as percentage of total renal cells.

### Western immunoblotting

Neonatal kidneys from mice undergoing UUO surgery or sham operation collected at day 3, 7, 14 and 21 of life were homogenized in protein lysis buffer (Tris 50 mM, 2% SDS, 1mM Na_3_VO_4_) containing proteinase inhibitors (Complete Mini, Roche Diagnostics) and benzonase (Novagen, Merck) and centrifuged for 10 minutes at 16,000 x g. Protein content was measured using a BCA kit (Thermo Scientific). 20µg of protein of each sample was separated on polyacrylamide gels at 160V for 80 minutes and blotted onto PVDF-membranes (Millipore, 80mA/membrane, 90 minutes). After blocking for 2h in Tris-buffered saline with Tween-20 containing 5% nonfat dry milk and/or BSA, blots were incubated with primary antibodies 2h at room temperature or at 4°C overnight. Blots were washed and incubated with horseradish peroxidase-conjugated secondary antibody for 1h at room temperature. Immune complexes were detected using enhanced chemiluminescence. Blots were exposed to X-ray films (Kodak), and protein bands were quantified using densitometry. Each band represents one single mouse kidney.

### Immunohistochemistry of human kidney section

Sections of paraffin-embedded kidney biopsies (1-2µm) were stained with standard periodic acid Schiff reagent (PAS), chloroacetatesterase (CAE) or anti-human UMOD antibody.

### Statistics

Data are presented as mean±SEM, cumulative frequency or representative images as depicted in the figure legends. Group sizes were chosen based on previous experiments. GraphPad Prism 7 software (GraphPad Software Inc.) was used to analyze data and to illustrate graphs. Statistical tests were carried out according to the number of groups being compared. For pairwise comparison, an unpaired student’s t-test and for more than two experimental groups, either a one-way or a two-way analysis of variance (ANOVA) with either a Tukey’s or a Sidak’s *post-hoc* test was performed. P-values <0.05 were considered statistically significant and indicated as follows: *: <0.05, **: <0.01, ***: <0.005.

### Study approval

The government of Oberbayern, Germany, approved all animal experiments, AZ 55.2-1-54-2532-76-12-2012 and ROB55.2-2532.Vet_02-17-102 and AZ 55.2-1-54-2532-118-11. Blood sampling from healthy volunteers was approved by the ethic committee from the LMU München (Az. 611-15).

## AUTHOR CONTRIBUTION

RI and BL-S designed and conducted experiments, analyzed data and wrote the manuscript. TS, HB, GH, FP, BP and BU acquired and analyzed data. HM, CAR, VV and JS provided their expertise and critical reagents. MS and MP designed experiments and wrote the manuscript.

## ACKNOWLEGMENTS

We thank Olivier Devuyst for critical discussion. In addition, we thank Susanne Bierschenk, Nadine Schmidt, Dorothee Gössel, and Ursula Keller for excellent technical assistance as well as the core facility BioImaging at the Biomedical Center, LMU Planegg-Martinsried, Germany for their help with microscopy.

This work was supported by the German Research Foundation (DFG) collaborative research grant SFB914, projects B01 (MS), B03 (CAR), and B11 (MP), DFG La 1257/5-1 (BL-S) and by the FöFoLe-Program of the Medical Faculty, LMU Munich (MP and MS).

## Conflict of interest statement

The authors have declared that no conflict of interest exists.

## Notes

### Competing Interest Statement

The authors have declared no competing interest.

