## Supplementary material for "Extratubular polymerized uromodulin induces leukocyte recruitment and inflammation *in vivo*": Suplemental Material

Supplemental Figure 1

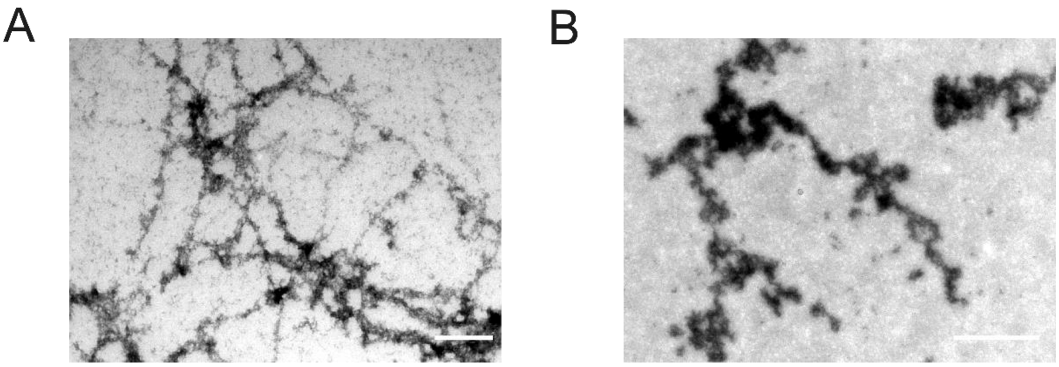


**Supplemental Figure 1.** (A) Representative transmission electron micrograph display (A) a filamentous network of human polymerized UMOD and (B) aggregates of recombinant murine UMOD (scale bars=1µm).

### Supplemental Figure 2

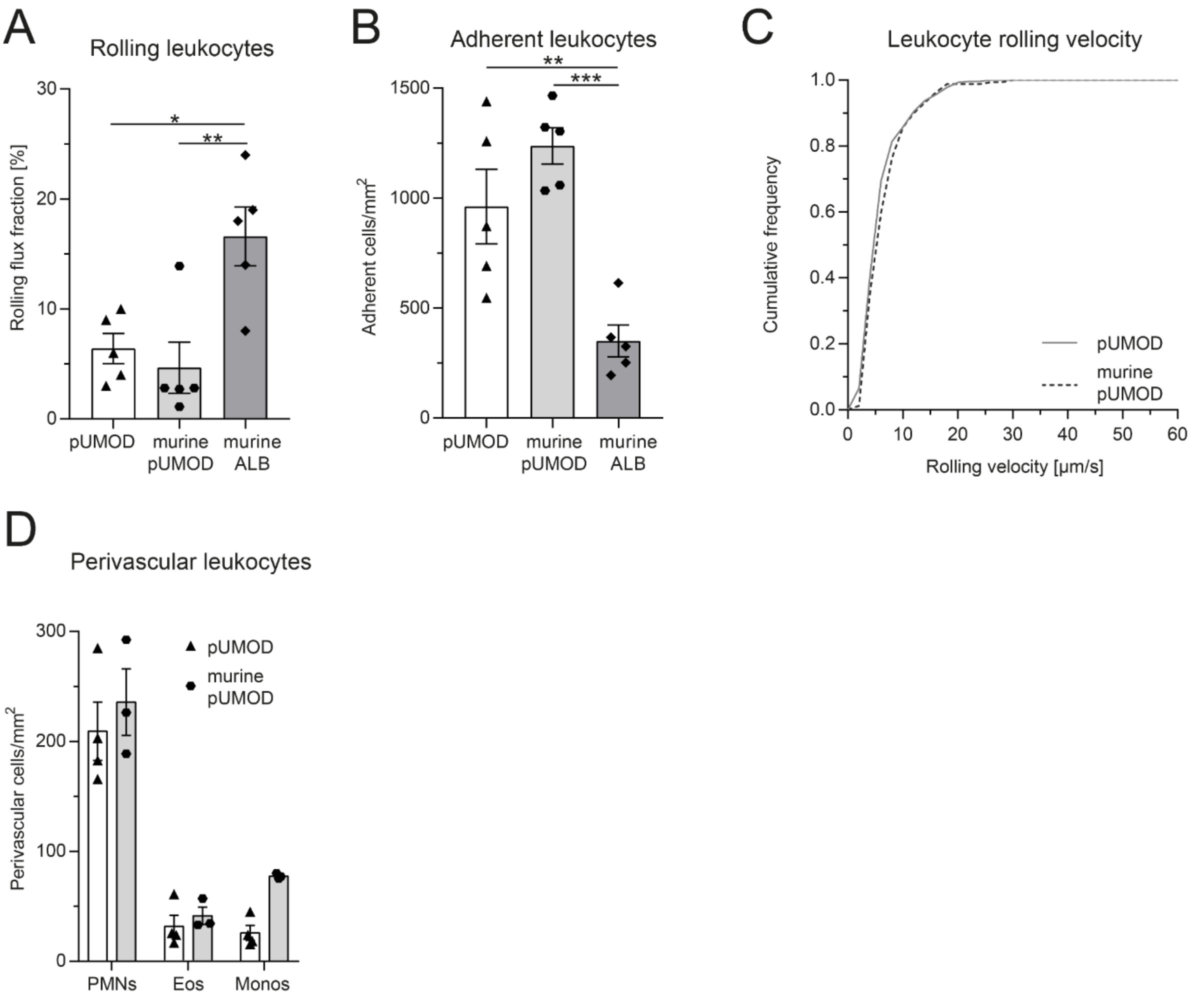


**Supplemental Figure 2 Intrascrotal application of murine polymerized UMOD (murine pUMOD) induces acute inflammation in postcapillary venules of the mouse cremaster muscle similar to pUMOD *in vivo*.** pUMOD, murine pUMOD or mouse albumin (murine ALB) were injected i.s. two hours prior to intravital microscopy. (**A**) Leukocyte rolling flux fraction and (**B**) number of adherent leukocytes were assessed in 23 (pUMOD, same values as in Figure 1A), 19 (murine pUMOD) and 29 (murine ALB) postcapillary venules of the mouse cremaster in n=5 mice per group (one-way ANOVA, Tukey's multiple comparison). (**C**) Leukocyte rolling velocities of n=274 (pUMOD, same values as in Figure 1C) and 161 (murine pUMOD) cells in 5 mice per group were analyzed (unpaired sudents t-test). (**D**) Number of perivascular neutrophils (PMNs), eosinophils (Eos) and monocytes (Monos) were calculated in 17 (murine pUMOD) and 54 (pUMOD, same values as in Figure 1E) perivascular regions of n=3-4 mice (two-way ANOVA, Sidak's multiple comparison). Murine pUMOD affected the leukocyte recruitment cascade similar to pUMOD. *: $p \leq 0.05$, **: $p \leq 0.01$, ***: $p \leq 0.005$, data is presented as mean±SEM and cumulative frequency.

### Supplemental Figure 3

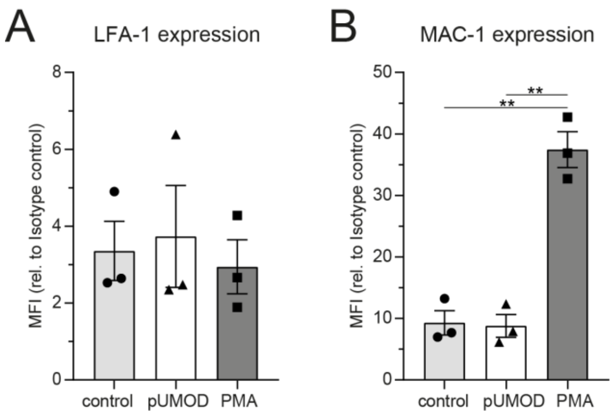


**Supplemental Figure 3 pUMOD does not alter surface expression levels of β2 integrins on human neutrophils.** Surface expression levels of (**A**) LFA-1 and (**B**) Mac-1 on isolated human neutrophils after stimulation with pUMOD, PMA and vehicle control were analyzed (n=3 independent experiments, one-way ANOVA, Tukey's multiple comparison). **: $p \leq 0.01$, data is presented as mean±SEM.

**Supplemental Table 1 Microvascular parameters.** Vessel diameter, centerline blood flow velocity, wall shear rate and white blood cell counts (WBC) of WT mice stimulated with vehicle control (control), polymerized UMOD (pUMOD) or human albumin (hALB) two hours prior to intravital microscopy (mean±SEM; 1-way ANOVA, Tukey's multiple comparison).

| | n (mice) | n (venules) | Diameter [µm] | Centerline velocity [$\mu m\ s^{-1}$] | Wall shear rate [$s^{-1}$] | WBC [$\mu l^{-1}$] |
|---|---|---|---|---|---|---|
| **control** | 4 | 17 | 31±1 | 1682±139 | 1297±86 | 4190±643 |
| **pUMOD** | 5 | 23 | 31±1 | 1926±159 | 1554±116 | 5858±1081 |
| **hALB** | 5 | 18 | 31±1 | 1917±148 | 1549±136 | 6342±490 |
| | | | ns. (p=0.8081) | ns. (p=0.4686) | ns. (p=0.2349) | ns. (p=0.8386) |

**Supplemental Table 2 Microvascular parameters.** Vessel diameter, centerline blood flow velocity, wall shear rate and white blood cell counts (WBC) of WT mice stimulated with pUMOD, murine pUMOD or murine ALB two hours prior to intravital microscopy (mean±SEM; 1-way ANOVA, Tukey's multiple comparison). pUMOD values are the same as in Supplemental Table 1.

| | **n (mice)** | **n (venules)** | **Diameter [µm]** | **Centerline velocity [µm $s^{-1}$]** | **Wall shear rate [$s^{-1}$]** | **WBC [$\mu l^{-1}$]** |
|---|---|---|---|---|---|---|
| **pUMOD** | 5 | 23 | 31±1 | 1926±159 | 1554±116 | 5858±1081 |
| **murine pUMOD** | 5 | 19 | 32±1 | 1974±168 | 1533±135 | 6788±572 |
| **murine ALB** | 5 | 29 | 31±1 | 1855±129 | 1466±106 | 3943±626 |
| | | | ns. (p=0.6631) | ns. (p=0.8512) | ns. (p=0.8458) | ns. (p=0.0690) |

**Supplemental Table 3 Microvascular parameters.** Vessel diameter, centerline blood flow velocity, wall shear rate and white blood cell counts (WBC) of WT mice stimulated with TNF-α, or a combination of TNF-α with pUMOD two hours prior to intravital microscopy (mean±SEM; 1-way ANOVA, Tukey's multiple comparison).

| | **n (mice)** | **n (venules)** | **Diameter [µm]** | **Centerline velocity [µm $s^{-1}$]** | **Wall shear rate [$s^{-1}$]** | **WBC [$\mu l^{-1}$]** |
|---|---|---|---|---|---|---|
| **TNF-α** | 5 | 34 | 26±1 | 1415±87 | 1353±9 | 4120±392 |
| **TNF-α +pUMOD** | 5 | 35 | 28±1 | 1446±75 | 1293±52 | 4900±694 |
| | | | ns. (p=0.4181) | ns. (p=0.7882) | ns. (p=0.5682) | ns. (p=0.3563) |
